# Cryo-FIB Lift-out and Electron Tomography Workflow for Bacteria-Nanopillar Interface Imaging Under Native Conditions: Investigating Dragonfly Inspired Bactericidal Titanium Surfaces

**DOI:** 10.1101/2025.11.19.688262

**Authors:** Chaturanga D. Bandara, Dominik Pinkas, Martina Zanova, Martin Uher, Judith Mantell, Bo Su, Angela H. Nobbs, Paul Verkade

## Abstract

Dragonfly- and cicada-wing-inspired titanium nanostructured surfaces exhibit promising bactericidal properties. However, direct visualisation of the bacteria–material interface under hydrated conditions remains limited, restricting experimental interrogation how bacteria interact with nanostructured surfaces. This limitation reflects a long-standing methodological gap, as visualisation of bacteria–nanostructure interfaces has largely relied upon indirect or dehydrated imaging approaches. Cryo-electron tomography (cryo-ET) enables three-dimensional visualisation of cellular ultrastructure in a native hydrated environment, but imaging bacteria attached to metallic nanostructures by cryo-ET requires preparation of thin electron-transparent lamellae. Obtaining such lamellae from vitrified bacteria on titanium substrates is technically challenging because cryo-focused ion beam (cryo-FIB) milling must simultaneously section soft biological material and hard metal whilst bacteria embedded in vitreous ice are not directly visible. Here, we establish a correlative cryogenic workflow enabling targeted extraction and cryo-ET imaging of defined bacterium–nanopillar interfaces. Individual bacteria interacting with titanium nanopillars are identified by correlative cryo-fluorescence microscopy, followed by targeted cryo-FIB lift-out, transfer to a receiver grid, and thinning into electron-transparent lamellae. The data presented demonstrate that bacteria–nanostructure interfaces can be targeted, extracted, and structurally analysed in situ under fully hydrated conditions. This workflow provides a methodological framework for future cryo-ET studies of bacteria–nanotopography interactions.

## Introduction

The rise of antimicrobial resistance (AMR) poses a major global health threat, driving the need for innovative strategies to prevent bacterial infection of medical implants. Biomimetic nanostructured surfaces with bactericidal properties have emerged as a promising solution in the last decade, particularly those inspired by natural nanopillars on dragonfly and cicada wings.^1–7^ Research has shown that bacterial cells that adhere to these biomimetic nanopillar surfaces become non-viable, highlighting a promising alternative to traditional chemical antimicrobials. Over the past decade, significant effort has been devoted to developing bioinspired nanostructured surfaces with enhanced bactericidal effects across a broad range of materials.^8–13^ Despite this progress, a comprehensive understanding of the bactericidal mechanisms by which these nature-inspired nanostructures mediate their effects remains lacking. Filling this research gap has been fundamentally constrained by a lack of suitable imaging methods capable of capturing the bacteria-material interface in its native hydrated state. Developing such capability is a prerequisite for obtaining meaningful mechanistic insight, and represents a major unmet methodological challenge in the field.

Several computational models have proposed different interactions at the bacteria-material interface such as membrane stretching, rupture, and cell wall penetration by nanopillars; killing bacteria however, experimental validation of these events remains limited. Conventional imaging techniques, including scanning electron microscopy (SEM), transmission electron microscopy (TEM), and fluorescence microscopy, have been employed to study these interactions at the interface^5, 6, 14–18^ but each comes with distinct limitations. SEM provides a detailed view of bacterial morphology on nanopillar surfaces but cannot capture the critical bacteria-surface interface where interactions occur, whereas this interface can be studied using FIB-SEM. Fluorescence microscopy reports on membrane integrity but lacks the spatial resolution to visualise nanopillars interacting with the bacterial cell wall, or to resolve nanoscale structural changes in the cell wall, hence conclusions made regarding cell wall damage are typically inferred indirectly. TEM offers high-resolution imaging but requires thin, electron-transparent samples, restricting the ability to observe the interactions of an entire bacterial cell with nanopillars. Much of the current ultrastructural evidence for bacteria-nanostructure interactions has relied on high-resolution imaging approaches performed on chemically fixed, dehydrated and resin-embedded samples. Hence, interface information available to date is derived from conditions far from the native hydrated state and could have introduced artefacts that can obscure the true bacteria-nanopillar interactions. Therefore, understanding bacteria-nanopillar interactions requires imaging methods that preserve the native hydrated state at the bacteria-material interface.

To overcome these challenges, correlative cryo-microscopy methods that enable the characterisation of bacteria-nanopillar interactions in their near-native hydrated state (in the presence of liquid, without chemical fixation and dehydration) are urgently needed. Cryo-electron tomography (cryo-ET) facilitates the investigation of ultrastructural studies of bacteria within their near-native state on nanostructured substrates such as titanium medical implants. Gaining insights into the ultrastructure of bacteria-material interfaces in relation to their surroundings is essential for connecting structure with function for biomimetic antibacterial surfaces. Given that most biological specimens are too thick for direct imaging using cryo-transmission electron microscope (cryo-TEM), techniques for reducing sample thickness below 200 nm is necessary for electron transparency^19, 20^ have been crucial for making cryoET of internal cell regions and bacteria-surface interfaces possible.

Here we present a correlative cryo-microscopy workflow integrating cryo-fluoresence, cryo-FIB liftout and cryo-TEM for characterisation of bacteria-nanopillar interactions on dragonfly wing-inspired titanium nanostructured surfaces, in a near-native, fully hydrated state. This approach directly eliminates the fixation and dehydration requirements inherent to conventional EM methods used in our previous studies of these interactions,^5, 6, 14, 21^, and provides new opportunities to contribute native-state interfacial data that can complement computational models and better inform the design optimisation of nature-inspired antibacterial surfaces for medical implants.

Our correlative cryo-microscopy workflow integrates complementary imaging and micromachining techniques all under cryogenic conditions. Each step is designed to overcome a specific limitation of conventional approaches, and enables targeted interrogation of the bacteria-nanopillar interface under hydrated conditions. Cryo-vitrification preserves bacteria in a fully hydrated, chemically unperturbed state by rapidly immobilising the surrounding liquid, eliminating the need for chemical fixation, dehydration, or resin embedding. Cryo-fluorescence microscopy then localises individual bacteria on nanostructured surfaces within the vitrified sample — a critical step for targeted downstream processing, as bacteria are not directly visible in cryo-FIB for milling. Cryo-FIB lift-out extracts a defined volume containing the bacteria–nanopillar interface, which is then thinned into an electron-transparent lamella maintained at cryo temperature. Finally, cryo-ET visualises the interface at nanometre resolution under fully hydrated conditions, without the heavy-metal staining required by conventional TEM. Together, this cryo correlative pipeline enables targeted structural interrogation of bacteria–nanostructure interactions that were previously inaccessible, providing a methodological framework for future studies of bacterial interactions with bioinspired antimicrobial surfaces.

## Materials and Methods

### Fabrication of nanopillars on titanium by thermal oxidation

Grade V titanium alloy (Ti-6Al-4V) sheets with 75 µm thickness were laser cut (Pharos PH1-10 laser operated a 1030nm. Output 10W.) into circles of 3 mm or 1.4 mm in diameter. Then these were hand polished using silicon carbide grit sizes (#2000 and #4000). Then, to remove surface contaminants, titanium disks were placed in an ultrasonic bath (Grant Scientific UXB series), first in acetone followed by distilled water at 40 °C for 15 minutes each using 100% power. Then samples were placed into ethanol (99.9%) for 10 minutes before air drying. Thermal oxidation was conducted as previously reported ^14, 21,8, 15^ Briefly, titanium samples were sealed inside a horizontal alumina tube (120 cm x 11 cm outer x 9 cm inner) positioned in the tube furnace (Elite Thermal Systems Ltd). Prior to thermal oxidation, the furnace was purged with inert argon gas (Ar) for 30 minutes with 220 sccm to remove oxygen and achieve a one-directional flow. Following the purging, heating started at 10 °C per minute until 850 °C. While maintaining the temperature, Ar was redirected via a sealed bubbler with acetone (99.9%), maintained at 25 °C for 20 minutes. Then the purging via acetone was stopped and Ar was redirected to the tube and cooling was started. This resulted in a black colouration of the samples (**Figure 1a**). After cooling to room temperature, heating was reinitiated after airing the chamber with ambient air, heated to 600 °C and maintained for 30 minutes for annealing, and then samples were allowed to cool to room temperature overnight. This served to remove the carbon from the samples (Figure 1b). These samples served as the substrates for our experiments.

**Figure 1:**
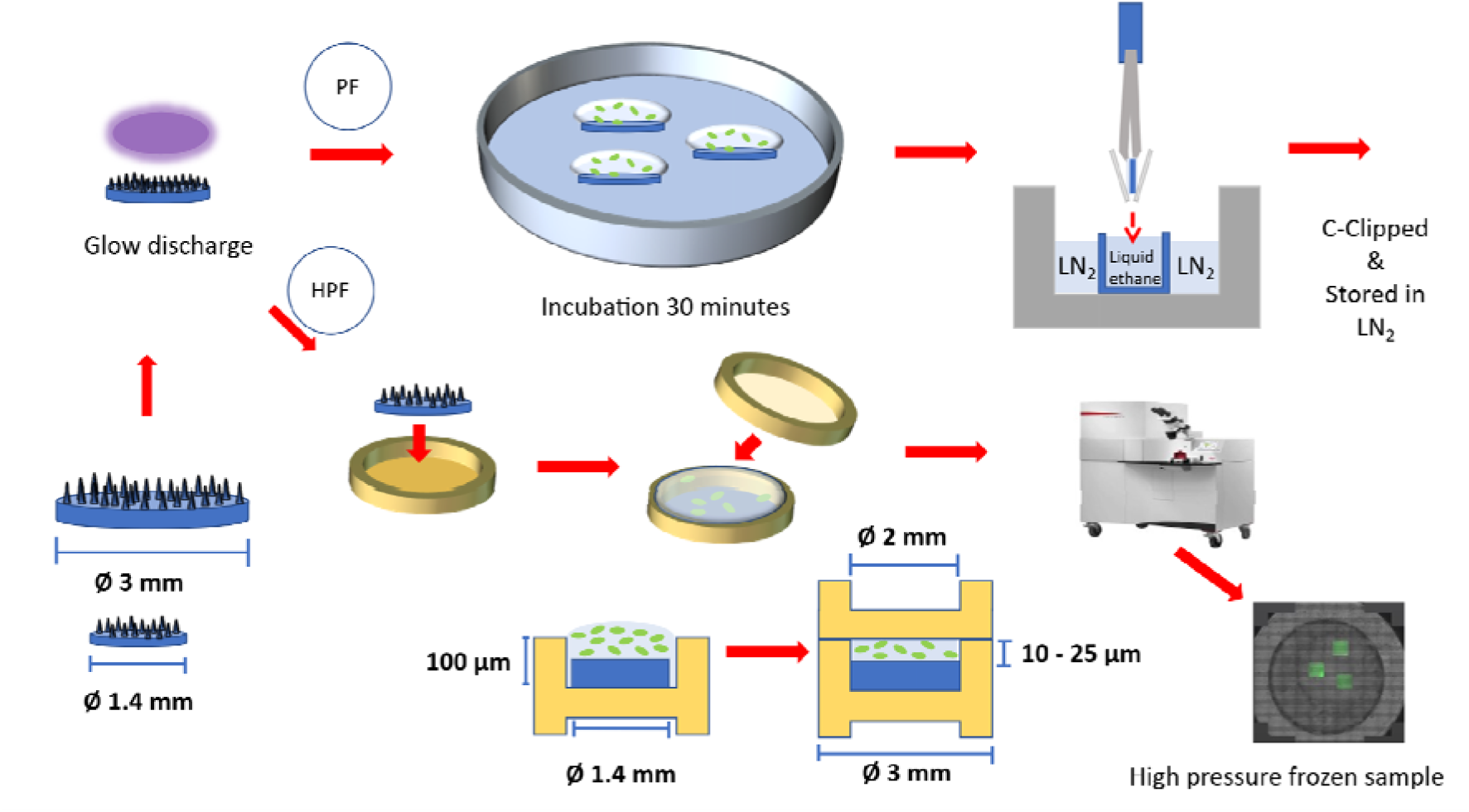
Illustration of cryo vitrification of samples. PF: Plunge freezing, HPF: High pressure freezing. Cleaned substrates are glow discharged, inoculated for 30 minutes, vitrified either with PF or HPF, and then stored in liquid nitrogen until further analysis. For further details, refer to text.

### Bacterial culture

*Escherichia coli* strain K12 was used as a model bacterium in this study. Luria Bertani broth cultures were incubated for 16 h at 37 °C, 200 rpm, then subcultured to OD_600_ 0.1 and grown to mid-exponential phase before use in the experiment. For incubation on titanium substrates, cultures were diluted to OD_600_ 0.1 in Luria Bertani broth. Titanium samples were sterilised in 70% ethanol, washed in dH_2_O, dried and stored until the experiment. Immediately prior to inoculation, titanium substrates were glow-discharged using Leica EM ACE600 (Leica Microsystems) to become hydrophilic. Without this step, inoculation was challenging, as the titanium sample bound to the end of the pipette tip when liquid transfer.

### Overview of cryogenic imaging workflow

To preserve the native ultrastructure and spatial context of bacteria at the nanopillar interface, imaging was performed under fully hydrated conditions using cryo-FIB-SEM, cryo-fluorescence and cryo-TEM. This approach enabled high-resolution visualisation of bacterial cells on biomimetic nanostructured surfaces without the chemical fixation, dehydration, or heavy metal staining required for conventional EM preparation. To maintain *in situ* positioning, bacteria were vitrified directly on titanium test samples, ensuring that cells remained undisturbed at their original locations. The cryogenic workflow preserved the physiological integrity of both the cells and the surrounding material, allowing accurate correlation between morphology and function in a near-native hydrated environment at the time of imaging.

Achieving effective vitrification directly on test substrates was a critical challenge. Both plunge freezing and HPF methods were explored in the experiments. Plunge freezing yielded a vitrified layer ∼5 µm thick, while HPF resulted in a thicker layer of ∼25 µm. As bacterial cells (∼1 µm in diameter) interacting with the surface were entirely embedded within the vitrified volume, they were not visible by SEM alone, complicating direct targeting of bacteria for cryo-ET.

To address this, correlative cryo-fluorescence microscopy was employed to localise regions of interest prior to FIB milling. However, the absence of fiducial features that were simultaneously visible in both fluorescence and electron modalities limited the precision of image registration. This was mitigated by two approaches; 1) by adapting to surface cracks or 2) making fiducial marks on the vitrified sample surface using FIB-SEM. A final constraint was the requirement for electron-transparent lamellae for cryo-TEM. To achieve this, selected regions identified by cryo-fluorescence were isolated and extracted onto a TEM grid using FIB-SEM with a cryo-cooled nanomanipulator and finally thinned to <200 nm, enabling high-resolution structural analysis of the bacteria-nanopillar interface by cryo-TEM.

In the following sections, different methods and challenges faced and how they were mitigated are extensively discussed.

### Cryo vitrification

Cryo-lift-out starts with the cryo vitrification step to achieve a vitreous ice sample. We tested both plunge freezing (PF) and high pressure freezing (HPF) in this experiment, as illustrated in Figure 1.

#### Plunge freezing

For plunge freezing, cells were vitrified using a Leica EM GP2 (Leica Microsystems). A 20 µL aliquot of bacterial suspension was applied to the sample and incubated for 20 minutes at room temperature. Then, 10 µL of SYTO9 was added to stain the genetic material of the bacterial cells. After 10 minutes, these samples were transferred to a Leica EM GP2 plunge freezer at humidity 80%, samples were blotted one-sided for 4 seconds with filter paper and immediately frozen in liquid ethane/propane mixture maintained at liquid nitrogen temperature (Figure 1). Samples were then clipped into a TEM AutoGrid (Nanosoft) cartridge with the cell side facing down and stored in liquid nitrogen until imaging.

#### High-pressure freezing

High-pressure freezing was performed using a Leica EM ICE system. Samples at room temperature were added with 50 µL of *E. coli* suspension for 20 minutes in 3 mm copper planchets. Syto9 (50 µL) was then added for a further 10 minutes to stain the bacterial cells. These titanium samples with surface-attached bacteria were topped with a marked cap wetted with 1-hexadecene to facilitate detachment (Figure 1). Samples were rapidly vitrified under a pressure of 2100 bar and cooling to −196 °C within milliseconds in the Leica EM ICE high pressure freezer. The resulting vitrified media layer was approximately 25 µm thick (see Figure 1), fully embedding the bacterial cells and preserving their hydrated state. Following freezing, samples were stored in liquid nitrogen until further processing. The vitrified HPF carriers were manually opened within the cryo-CLEM loading station (Leica Microsystems, Vienna, Austria) immersed in liquid nitrogen. This was achieved by carefully pushing the planchets apart using tweezers. Once opened, the bottom carrier containing the sample was loaded into a custom-designed cartridge for transfer into the microscope.

#### Cryo-sample handling and transfer between different modalities

Maintaining vitrified samples in a stable cryogenic and contamination-free state throughout the correlative cryo-microscopy workflow is essential for preserving ultrastructure and enabling accurate multi-modal imaging of bacteria-nanopillar interfaces. A major logistical challenge in this workflow was the safe and efficient transfer of frozen-hydrated samples between multiple instruments, including the cryo-fluorescence microscope, cryo-coating system, cryo-FIB-SEM, and cryo-TEM, at different locations (IMG and TESCAN), whilst preventing devitrification and minimising ice contamination.

To address this, all sample handling and transfers were conducted using the Leica EM VCM system and Leica EM VCT500 shuttle inside the laboratory. The Leica EM VCM provided a temperature-controlled cryo-loading interface, maintained at liquid nitrogen temperature, allowing sample manipulation and orientation in a protective nitrogen atmosphere without devitrification and minimal frost contamination to exchange samples between the cryo-fluorescence microscope and cryo-FIB-SEM. For sample transportation between different locations, a dry shipper (CX-100) was used.

### Cryo light microscopy (cryo-LM)

The carrier containing the sample was loaded into a custom-designed cartridge for transfer into the Leica EM Cryo CLEM Thunder Imager system (Leica Microsystems, Austria). The sample was maintained in overpressure of cold nitrogen gas to maintain a vitrified state and avoid contamination with atmospheric humidity. Images were acquired with a 50x dry objective with NA of 0.9 (HC PL APO, CRYO CLEM, Leica Microsystems) equipped with a Leica DFC9000 GT camera. Light microscopy images were captured in reflected light and fluorescence (EX 450-490nm, EM 500-550nm) modes. Large areas were imaged via stitching of multiple individual exposures using the Leica Navigator module within the LAS-X software package. Images were captured at a frame size of 2048 x 2048 pixels, resulting in a pixel size of 0.13 µm. Manufacturer recommended system-optimized Z-steps of 0.44 µm were used. An exposure time of 100 ms in channel 1 (SYTO9, green fluorescence) and 1 ms in channel 2 (REF, reflection, grey) was used. To improve the clarity of the fluorescence datasets and enable reliable correlation with the cryo-FIB-SEM step, the Z-stacks were processed using the proprietary GPU accelerated Leica Thunder deconvolution technique in the LAS X software (Leica Microsystems, Vienna, Austria). The marked fiducials and fluorescence data were combined in Fiji, and the resulting composite images were exported in TIFF format and subsequently used as reference maps for cross-modality alignment.

### Cryo-correlation of fluorescence data and ion beam field of view for selecting a single-bacterium for lift-out

Since bacteria were hidden below the surface, and surface topography of vitrified samples did not indicate the presence of bacteria, cryo-fluorescence and cryo-FIB-SEM data were required to produce targeted cryo-lamellae from vitrified samples. Moreover, as the plasma FIB-SEM system lacked an integrated cryo-fluorescence microscope, it was essential to predefine and mark regions of interest for lamellae in advance. Mapping the entire sample at nanometer precision is impractical and prone to error due to the image stitching process, ice contamination, and the inherent complexity of spatial correlation. To overcome this challenge, we tested two different approaches to enable reliable localisation of bacterial targets with high spatial precision: first, using dominant surface features, then second, using marked fiducials via the Ga-FIB on the surface, as illustrated in **Figure 2**.

**Figure 2:**
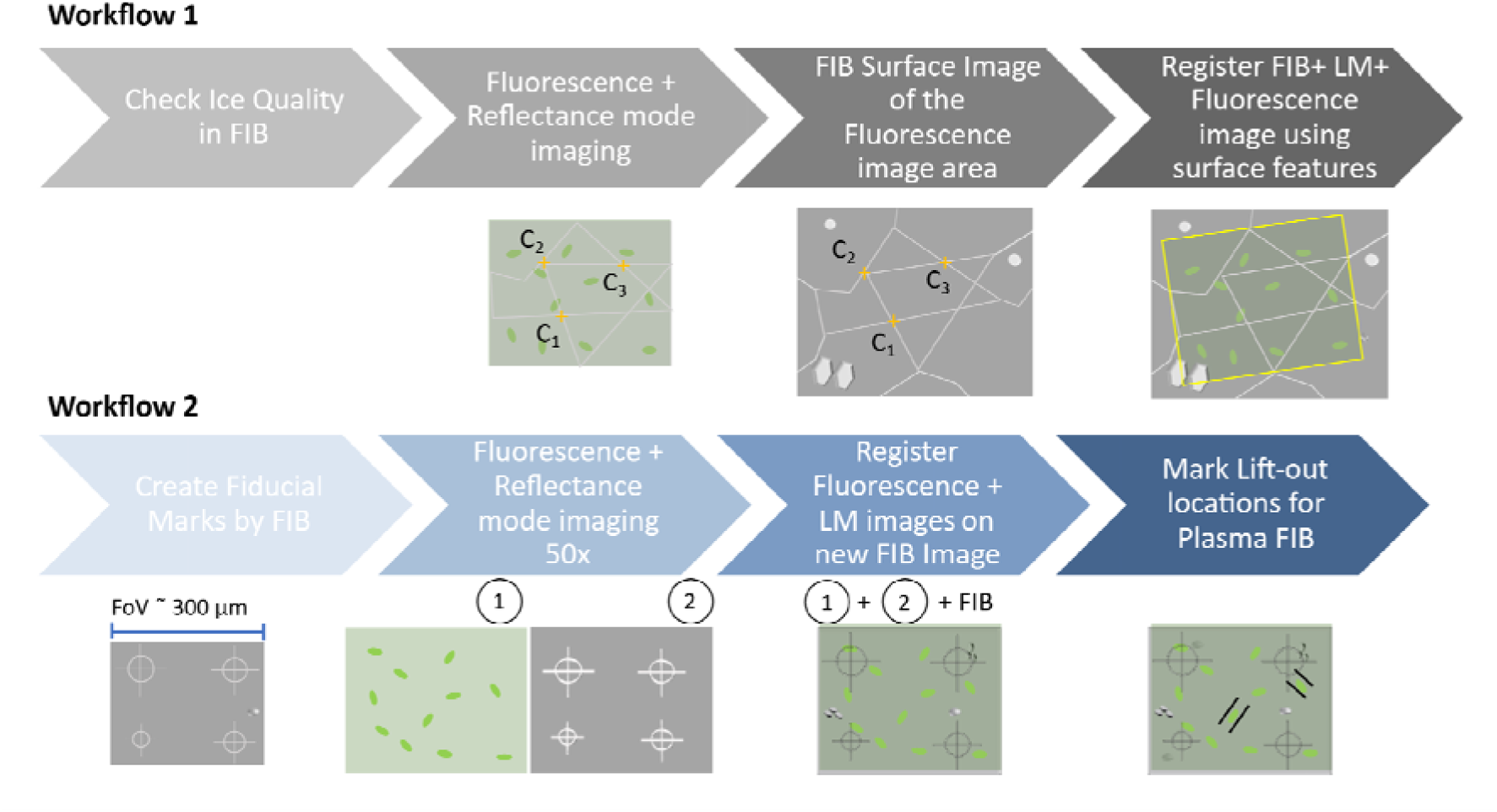
Illustration of two different correlative microscopy approaches used in this work. **Workflow 1**: With Tescan Essence software using surface cracks. **Workflow 2**: With defined fiducials on the surface of the sample using Ga-ion beam. Marked fiducials are visible in the 50x field of view in the reflectance mode. Their asymmetry helps to identify the sample when the fluorescence microscope provides flipped images. Finally, the lift-out locations are marked by two lines centring a single bacterium.

### Correlative Registration Protocol Based on Surface Features

A clearly recognisable feature on the sample surface was recognised using the fluorescence microscope. In this instance, a 10x10 µm FIB milled area surrounded with surface cracks was picked (**Figure 3** a,b). Then a reflectance light image and a fluorescence image of the same area was acquired and the channels were merged into a composite image. Surface features, including cracks, and ice forms were visible in the composite image, along with fluorescently labelled bacteria. Then the sample was transferred to the Ga-FIB instrument under cryo conditions, and the same field of view was relocated. Excessive ice formations were milled away by scanning with the ion beam at a lower magnification to facilitate recognition of the surface features. Then the reflected light image was overlayed using Tescan Coral software by selecting correlated positions (C1, C2, and C3 in Figure 2) in both FIB and imported reflected light image (Figure 3d). Then the acquired Z-stack was analysed to identify bacterial cells located at the bottom of the fluorescence volume (i.e., adjacent to the titanium nanopillar surface). Only positions of these surface-associated cells were selected for lamella preparation to enable investigation of bacteria-material interactions via cryo-ET (Figure 3e).

**Figure 3:**
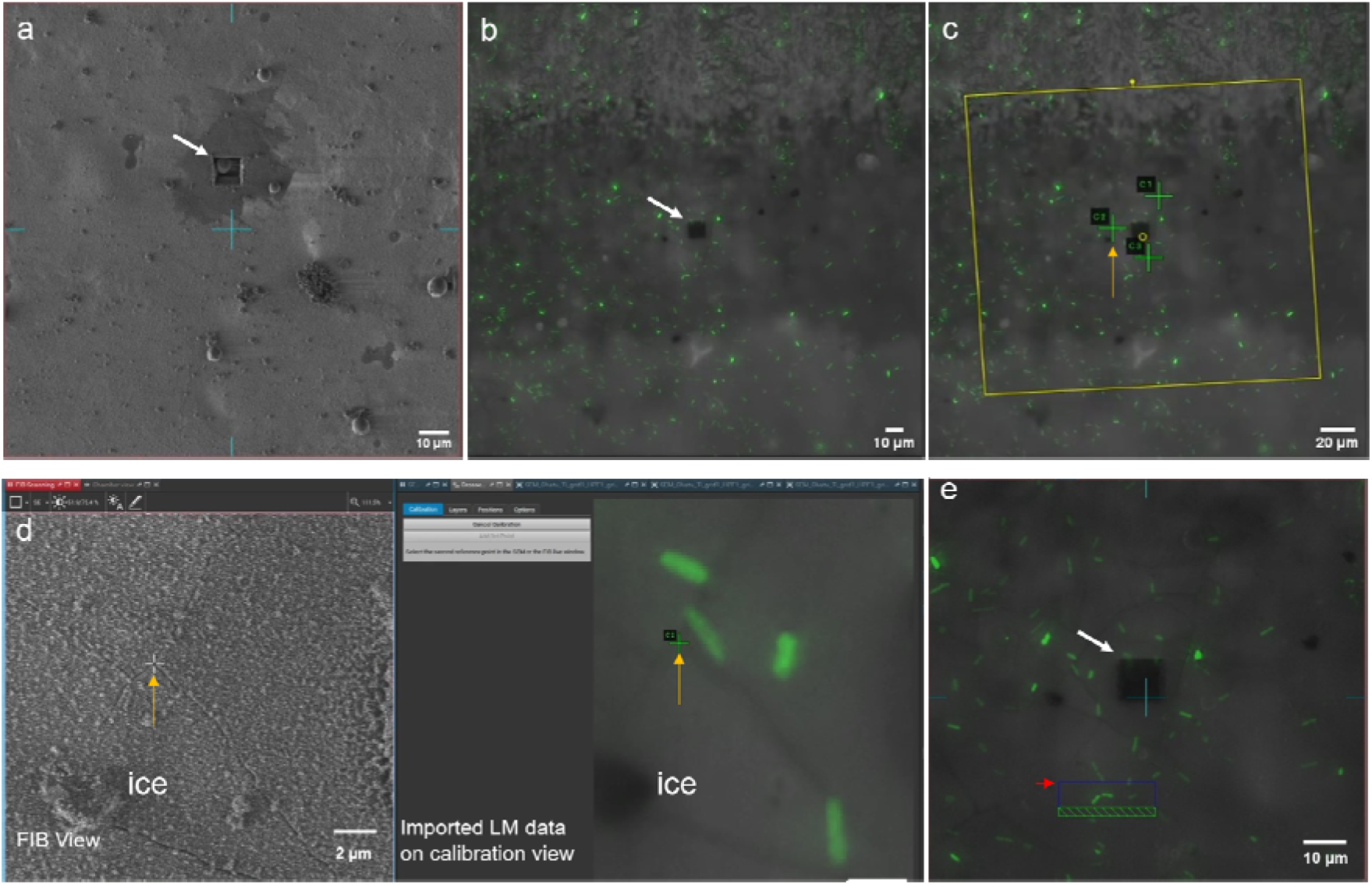
Correlation of fluorescence data with FIB-SEM image using the Tescan Essance software. a) FIB-SEM image showing the area where ice quality was checked (white arrow). This prominent FIB-milled square is clearly visible in both FIB-SEM and in light microscopy data during image correlation process. b) Overlayed brightfield and fluorescence data where the prominent FIB-milled area is recognised at 50x magnification. c). Successful registration of fluorescence data with FIB view. Three green crosses correspond to three alignment positions (C_1_-C_3_) used in the process. Yellow arrow indicates C2 position used in the process. d) Tescan Essence software interface showing FIB view (left window) and fluorescence image (right window) during selection of the second marker point during the image correlation process. The white and green crosses (indicated by yellow arrows) correspond to the alignment position C_2_. e) After image correlation, marking the position of two bacterial cells of interest in the FIB. Marked blue square is the slab area (red arrow) and green marking is the FIB marking site. Screen views are adjusted for brightness and contrast for clear visualisation.

### Fiducial Marker Fabrication and Correlative Registration Protocol

#### Fiducial marker creation

Three discrete regions on the vitrified sample were randomly selected and marked using the Ga-FIB Cryo-SEM system (**Figure 4a**). In each region, four fiducial patterns comprising crosses and circles of different sizes were milled in a square configuration to form a recognisable layout (Figure 2 workflow 2) visible under the cryo-fluorescence microscope with a 50x objective and fitting all fiducials within a single field of view of 300 µm wide (Figure 4).

**Figure 4:**
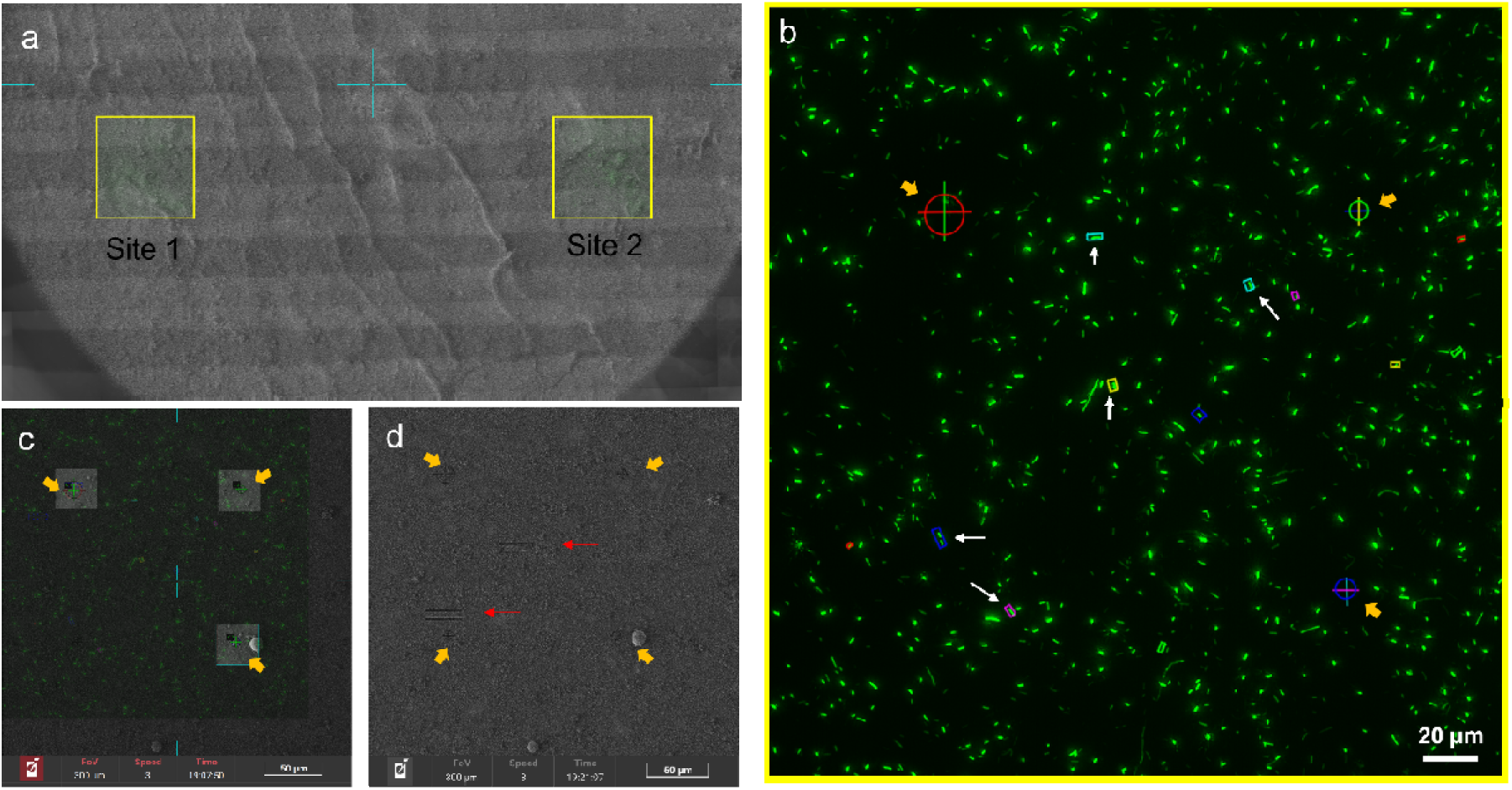
Fiducial marker-based identification of ROIs with bacteria for lift-out preparation. a) Brightfield image showing an overview of larger area of sample with two areas marked with fiducials (yellow squares). Only the bottom half of the 6 mm sample is shown here. Marked fiducials are not visible at this scale, but approximate locations of RoIs are identifiable. b) Fluorescence image of a single site shows bacteria in green and the fiducials visible in reflectance mode are marked on the fluorescence image (yellow arrows). Bacterial cells at the bottom of the Z stack are identified as regions of interest for FIB lift-outs. Several of these are marked in rectangles (white arrows). c) Fluorescence data is registered to FIB view and d) two lift-out positions per area are marked (red arrows) to perform lift-out without the need of a fluorescence microscope in the other facility.

#### Cryo-fluorescence imaging and target selection

In the cryo-fluorescence microscope, fiducial patterns were first identified using the reflected light channel. Each region of interest was centred within the square formed by the four fiducials (Figure 4b). A single reflectance mode image and fluorescence Z-stacks were acquired to image bacteria within the marked field. Then maximum intensity projections were generated, and fluorescence and reflected light channels were merged into a composite image. Z-stacks were analysed to identify bacterial cells located at the bottom of the fluorescence volume (i.e., adjacent to the titanium nanopillar surface). Only positions of these surface associated cells were selected for lamella preparation to enable investigation of bacteria-material interactions.

#### Correlative registration in cryo-FIB-SEM

Subsequently, the samples were transferred to the cryo-FIB-SEM for the second time, and the marked fiducials were located. Then the composite fluorescence data were imported into the Tescan CORAL module embedded within the Tescan ESSENCE software package used to control the FIB-SEM instrument. The fluorescence images were registered to the ion beam field of view using fiducial markers as anchor points (Figure 4c). Based on Z-stack data, precise lift-out locations were defined and targeted for lamella preparation. Identified lamella locations consisted of two 10 µm × 2 µm rectangles straddling a bacterium (Figure 4d). To mitigate the risk of fiducials being obscured by ice contamination during storage or transfer, several marked areas were generated as backups. Then Multi-scale “Zoom-in” maps of each fiducial-marked region were created in the Ga-FIB-SEM to guide the plasma FIB-SEM operator in efficiently navigating to regions of interest and executing high-precision lift-outs. Samples were stored in liquid nitrogen and transported to the next site in a dry shipper until plasma-FIB-SEM preparation.

Correlation of a light microscopy volume image and EM field of view presents inherent challenges due to the significant differences in resolution and field of view between the two modalities. Cryo-sample preparation and cryo-plasma FIB-SEM imaging were conducted at two separate facilities. As the plasma FIB-SEM system lacked an integrated cryo-fluorescence microscope, it was essential to predefine and mark regions of interest for lamellae in advance. Mapping the entire sample at nanometer precision is impractical and prone to error due to the image stitching process, ice contamination, and the inherent complexity of spatial correlation. To overcome this challenge, we developed a novel fiducial marker strategy to enable reliable localisation of bacterial targets with high spatial precision (**Figure 3&4**).

### Cryo FIB-SEM lift-out preparation

Cryo-FIB-SEM experiments were conducted on a Tescan AMBER Ga FIB-SEM and a Tescan AMBER X2 Xe plasma FIB-SEM system (Tescan), both equipped with Leica VCT500 cryo stages. In the Leica VCM loading station (Leica Micro-systems), HPF samples were mounted on a 30° pre-tilted prototype Leica cryo holder compatible with HPF carriers and Autogrids for lamella preparation. Using the Leica VCT500 Shuttle, the mounted sample holder was transferred into the Leica ACE600 cryo-sputter coater (Leica Microsystems) at a cryo-stage temperature of -150 °C and samples were sputter-coated with a 10 nm thick Pt layer. The samples were transferred to the Tescan FIB-SEM using the VCT500 Shuttle and a layer of platinum pre-cursor was deposited onto the sample surface to minimize the curtaining effect using gas injection system (GIS).^22^ The stage temperature was maintained below -150 °C and the system vacuum below 1.4 x10^-6^ mbar pressure. Due to the complexity and material properties of the titanium substrates used, a hybrid FIB strategy combining Ga-FIB and plasma FIB was adopted to optimise both milling efficiency and lamella quality.

#### 1. FIB-SEM - Ga

Vitrified samples were transferred under cryogenic temperatures into the Tescan AMBER Cryo Ga FIB-SEM (Tescan, Czech Republic) using a precooled cryo stage maintained below –150 °C. The preparation workflow consisted of four main steps: trench milling, rough polishing, slab extraction and final thinning.

Once the location of the bacterial cell was identified, three trenches were made around the target lamella area using the Ga ion beam, resulting in a block of material. Initially, stepwise cutting was performed with a depth of 30 μm. Then the redepositions were cleaned multiple times around the block of material and ‘*cleaning cross section*s’ were performed to make sure the desired level of depth was achieved. All sides were polished with a moderate current (30kV, 20nA) to make sure the target slab was separated from the bulk material from three sides. The sample was then rotated 180° and the slab was cut from the underside to separate it from the bulk material underneath yet still attached on one side. At this stage, the sample was stored overnight and on the next day, any ice contaminations were cleared by scanning with the ion beam at low magnification. After rotating the sample back to the initial milling direction, all sides were checked to ensure that the milled pillar was completely separated on three sides and the bottom, enabling the sample to proceed to the lift-out transfer step.

#### 2. FIB-SEM – Plasma

It was evident that Ga FIB was not efficient for extracting a titanium slab from bulk material, hence cryo Plasma FIB-SEM was adapted for lift-out preparation. Once vitrified, samples with marked lift-out locations were transferred under cryogenic temperatures into the dual-beam cryo-plasma FIB-SEM Tescan AMBER X2 (Tescan, Czech Republic) using a precooled cryo stage maintained below - 150 °C. The preparation workflow consisted of four main steps: trench milling, rough polishing, slab extraction and final thinning.

#### Trench milling and slab in-trench polishing using Xe Plasma FIB

Initially, pre-marked fiducials were identified in the SEM under low magnification (**Figure 5a**). A Pt layer was deposited using an ion beam assisted gas injection system to cover and protect the region of interest from ion damage during the milling process (Figure 5c). Using a xenon plasma ion beam at high current (30 kV, 60 nA), three trenches were milled around the target slab area, resulting in a thickness of 10 µm x 30 µm containing the pre-identified bacterium (Figure 5d-f). Then all slides were polished with a moderate current (30 kV, 20 nA) to reach a suitable slab size for cryo lift-out (Figure 5g). The sample was then rotated 180° and all the slabs were undercut to make sure that the ROI was fully separated from the bulk sample on the bottom side and yet hanging only from one end (Figure 5i). One slab was lifted out of the trench using Xe Plasma FIB. The two other slabs were transferred under cryo conditions back into storage until the lift-out and polishing steps (Figure 5).

**Figure 5:**
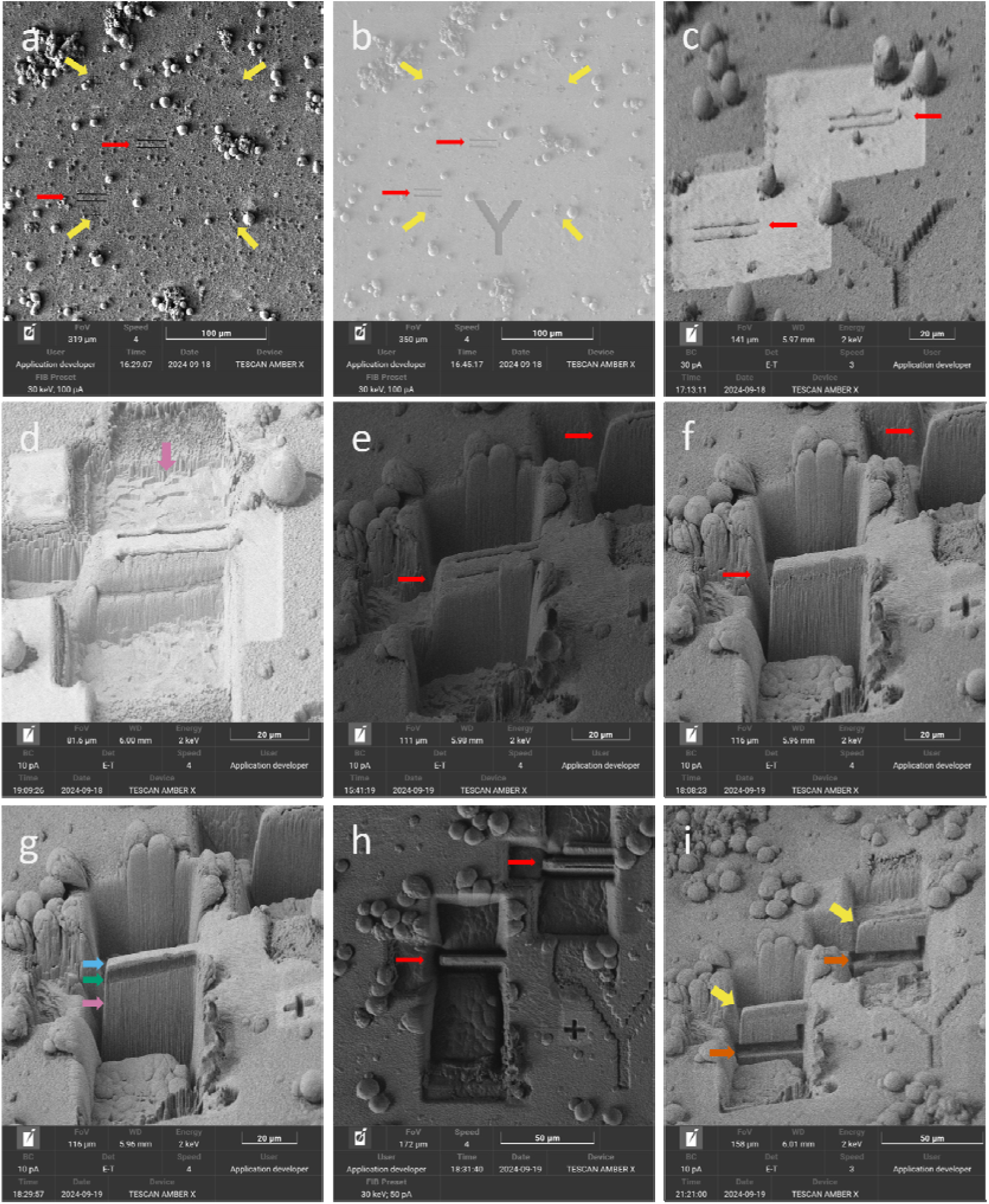
Isolation of a single bacterium for TEM tomography by targeted cryo-lift-out using FIB-SEM. a) The two yellow arrows point to two positions where two lamellae were to be prepared for cryo tomography. b) Letter Y was assigned and marked to this RoI before depositing a protective Pt layer. c) Area after Pt deposition, in preparation for milling. d) Trenches around one lamella being cut. Pink arrow indicates the exposed titanium. e) Trenches are cut to expose the areas of two lamellae. f) Deep trenches are cut, with the lamellae shown before polishing. g) One lamella is polished to show the different layers. Blue arrow, protective Pt layer; green arrow, vitrified water layer with bacteria; pink, titanium surface. h) Top view of the area in a) after trenches are cleared before the undercut. i) Undercut (orange arrow) has been performed, with the lamella now hanging to the titanium base from the side.

### Lift-out transfer

In the Ga-FIB-SEM, pre-milled areas were found under low magnification. The ice contamination from the transfer was removed using the Ga ions. A Tescan cryo-nanomanipulator (Tescan, Czech Republic) was used to lift out the slabs from the bulk sample (**Figure 6**). The nanomanipulator was brought closer to the slab and made 4 cuts of 200 nm size on the manipulator (30 kV, 3 nA). The milled material redeposited between the nanomanipulator and the slab attaching the slab to the manipulator (white arrow). To release the slab from the bulk sample, the remaining side connection on the bottom was cut off (30 kV, 3 nA) with an ion beam and the cryo slab was carefully lifted out (Figure 6b). The manipulator attached with the slab was brought closer to the FIB lift-out TEM grid, and the slab was attached to the TEM grid using redeposition (Figure 6c). Multiple cuts were made on the TEM grid very close to the slab to incur redeposition (yellow arrows). Then the slab was released from the nanomanipulator by sectioning the initial redeposition site (white arrow) (30 kV, 1 nA). These slabs were thinned from both sides using a stepwise reduction in ion current from 3 nA to 50 pA, until a final thickness suitable for cryoET of approximately 200 nm was achieved (Figure 6d-f).

**Figure 6:**
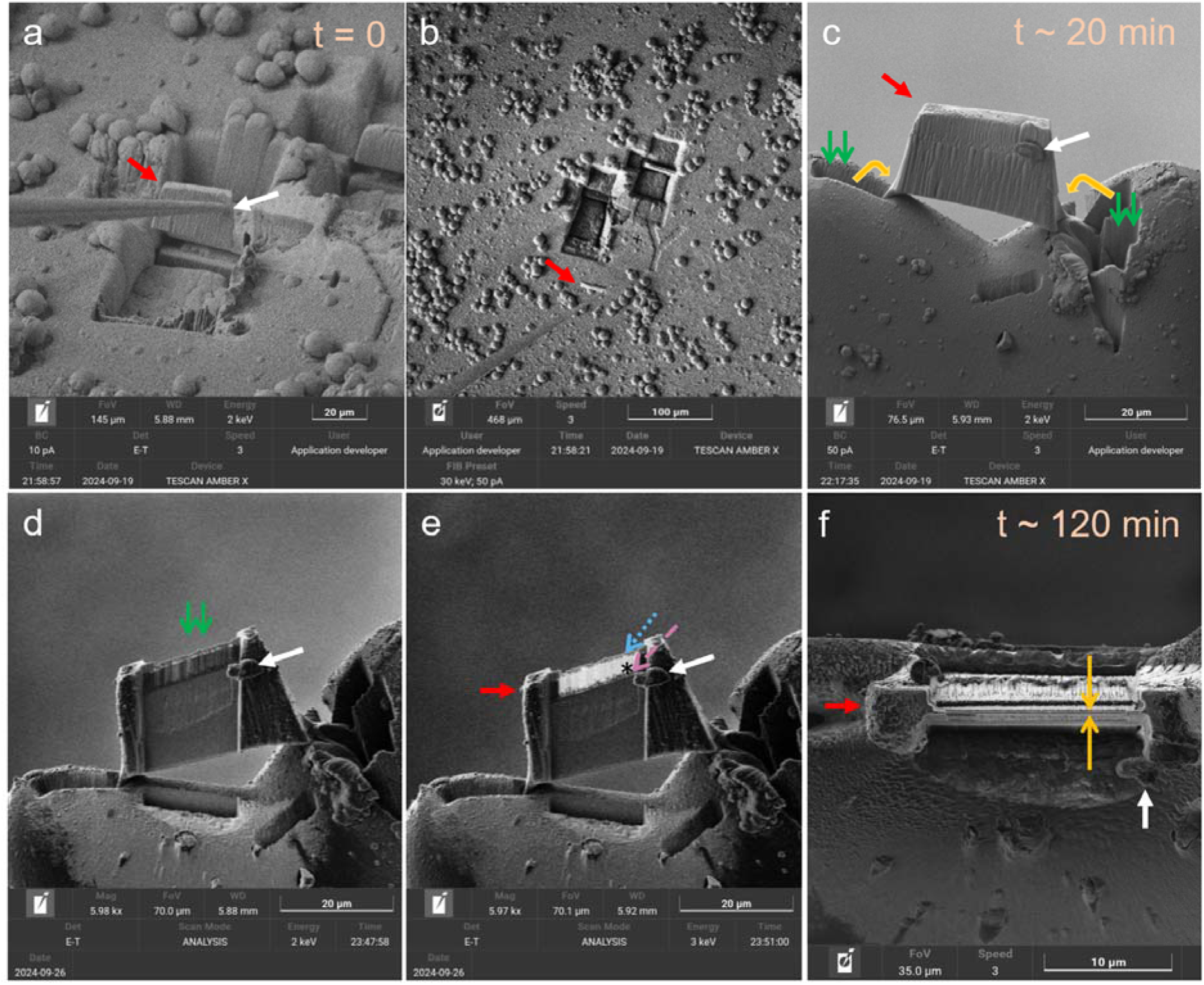
Lift-out of the slab on to a TEM grid to make an electron transparent lamella from the vitrified sample. a) The slab is detached from the bulk sample and lifted out using a cryo manipulator (red arrow points to the slab and white arrow points to the manipulator attachment). b) Top view image of the lamella being removed from the original site, while the second slab is still attached to the bulk sample. c) Attached lamella to TEM substrate. White arrow indicates the manipulator attached position, green arrows indicate the milling of Cu grid, yellow arrows indicate redeposition of Cu attaching the slab to the substrate, and d-e) progressive steps of lamella thinning and polishing to obtain electron transparent sample. When ready, different zones of lamella became visible: Pt deposition (dotted blue arrow), vitrified area on substrate (black asterix), nanopillars (dashed purple arrow). f) Top view after polishing of the lamella ready for TEM.

### Cryo-TEM

Thin lamellae were transferred into a cryo TEM (Jeol JEM-F200, Jeol, Japan) using a LN2 cooled Simple Origin Model 210 tomography cryo transfer holder (Simple origin, PA, USA). Samples were observed using a TVIPS TemCam XF-416 CMOS based camera (TVIPS, Germany). Tilt-series were acquired using the SerialEM software package from -60° to +60° tilt with 3° increment^23^, utilizing a dose symmetric acquisition scheme.^23, 24^

## Results

### Generation of nanopillars on titanium by thermal oxidation

The thermal oxidation method generated the nanopillar samples used for the study (**Figure 7**). After initial exposure to acetone, the sample appeared black in colour (Figure 7a) and after oxidation, samples appeared a yellow/blue colour (Figure 7b). Overall, the samples appeared flat (Figure 7c), but their surface comprised highly dense nanopillars (Figure 7d) and from the side view (Figure 7e), they were not flat.

**Figure 7:**
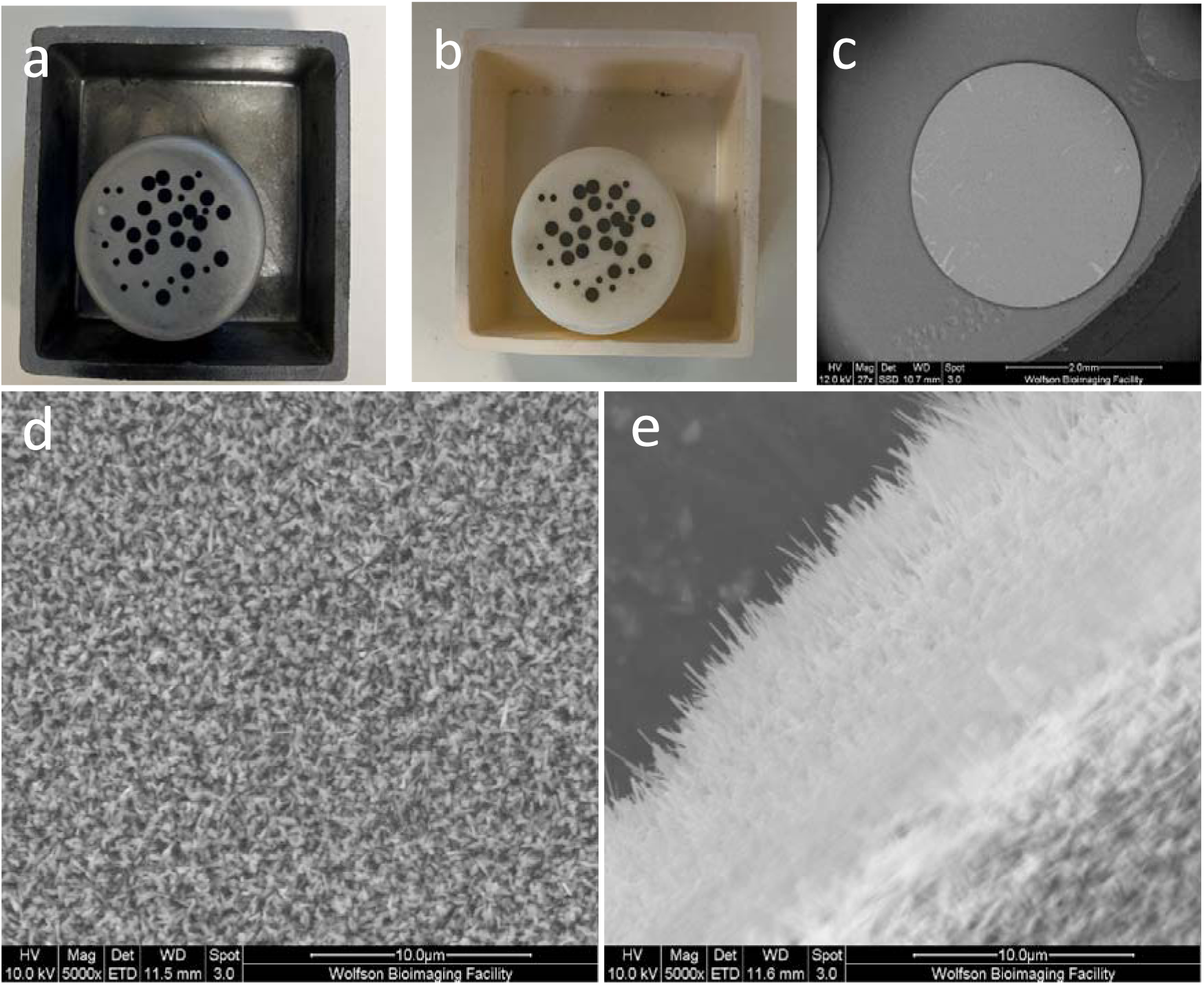
Fabricated nanopillar titanium sample. a) Samples after treated with acetone. b) Samples after oxidation in air. c) SEM micrograph showing the appearance of the nanopillar fabricated sample. d) SEM micrograph showing the top view of the fabricated surface with densely packed nanopillars. e) SEM micrograph showing a tilted view where protruding nanopillars are visible.

### Associated challenges

Targeting and isolating single bacteria and slabs (Figure 5,6) posed several difficulties. As shown in **Figure 8**, mainly Ga-FIB-SEM caused difficulties in lifting out the slab, as it was slower, and caused severe redeposition while milling the Ti substrate, compared to plasma-FIB. When redeposition occurred between the slab and the bulk material after the manipulator was attached and during the final step of separating the slab, the lift-out extraction was not efficient. Usually, this resulted in the manipulator detaching from the slab. On some occasions, poor attachment of the slab on the TEM grid had occurred (Figure 8b). Once the manipulator was detached, recovery of those slabs was unlikely. In the Ga-FIB, the milling rate was comparatively slow; hence, the sample had to be stored until the next session. Sometimes this caused ice contamination surrounding the slabs (**Figure 8c**). However, if this occurred after TEM lamella preparation, the sample was not usable.

**Figure 8:**
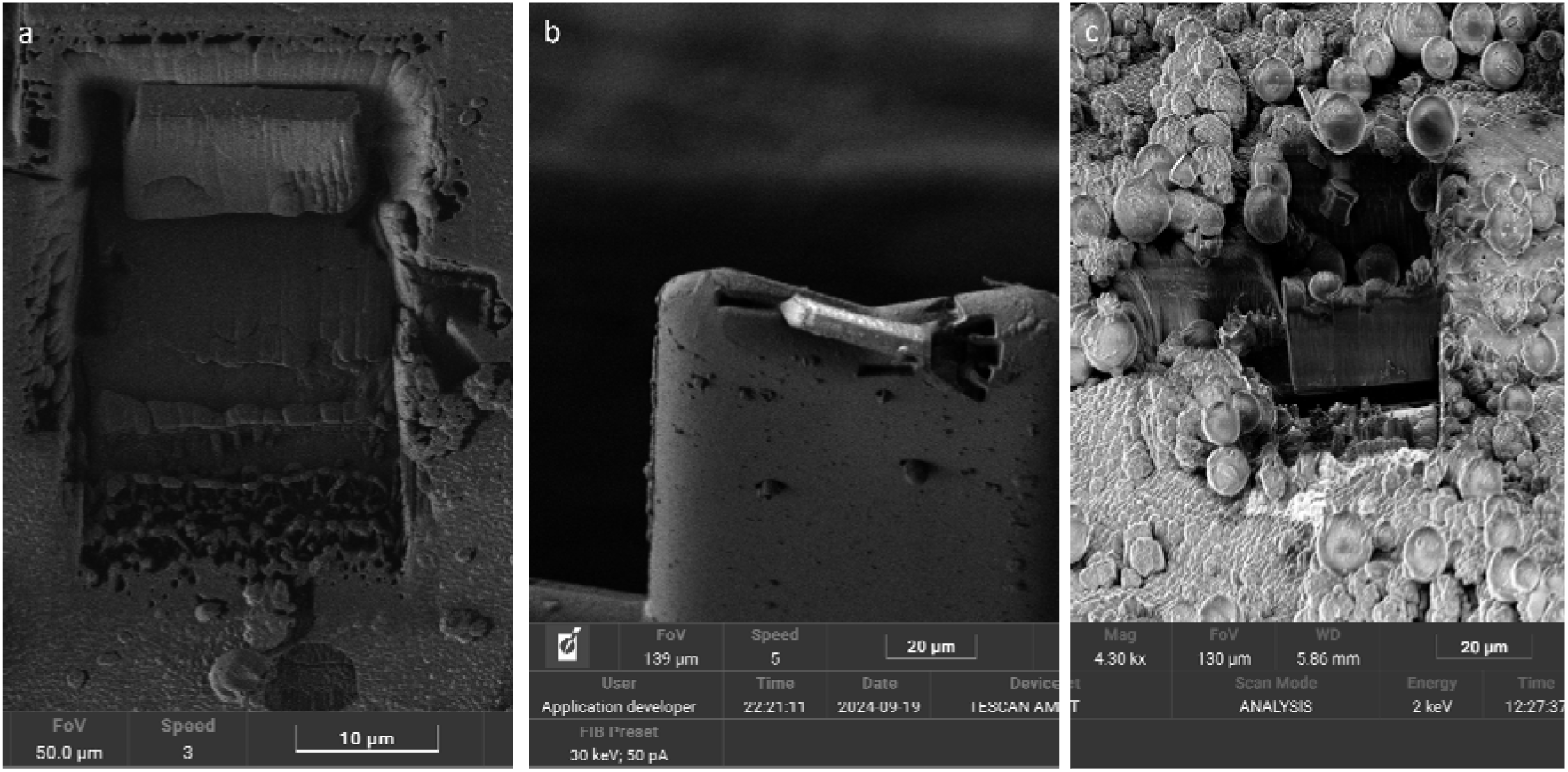
Challenges in the workflow. a) A lift-out was attached at the bottom during Ga-FIB-SEM preparation, hence it could not be lifted out easily. b) Lamella lift-out by Plasma FIB-SEM but failed upon transfer to the TEM grid. c) Ice contamination of the sample after storage and lamella is partially covered by ice.

### Visualisation of bacteria-nanopillar interface in hydrated state

Using our cryo-microscopy workflow, we have demonstrated that defined bacterium-nanopillar interfaces can be identified, targeted, and imaged in situ without chemical fixation, dehydration or resin embedding (**Figure 9**). The nanopillars and the bacterial cell wall are clearly resolved, and the interface region remains filled with the media without observable air gaps. A bacterium attached to the surface was intentionally targeted; however, a 100-200 nm separation is visible between the nanopillar tips and the bacterial cell wall. During initial fluorescence screening, nanopillars could not be resolved, precluding definitive identification of direct contact sites prior to FIB milling. This dataset nonetheless establishes that bacteria-nanostructured surface interfaces are accessible to cryo-microscopy investigation under native hydrated conditions, providing a methodological foundation for future studies aimed at mechanistic interrogation of bactericidal action on nanostructured surfaces.

**Figure 9:**
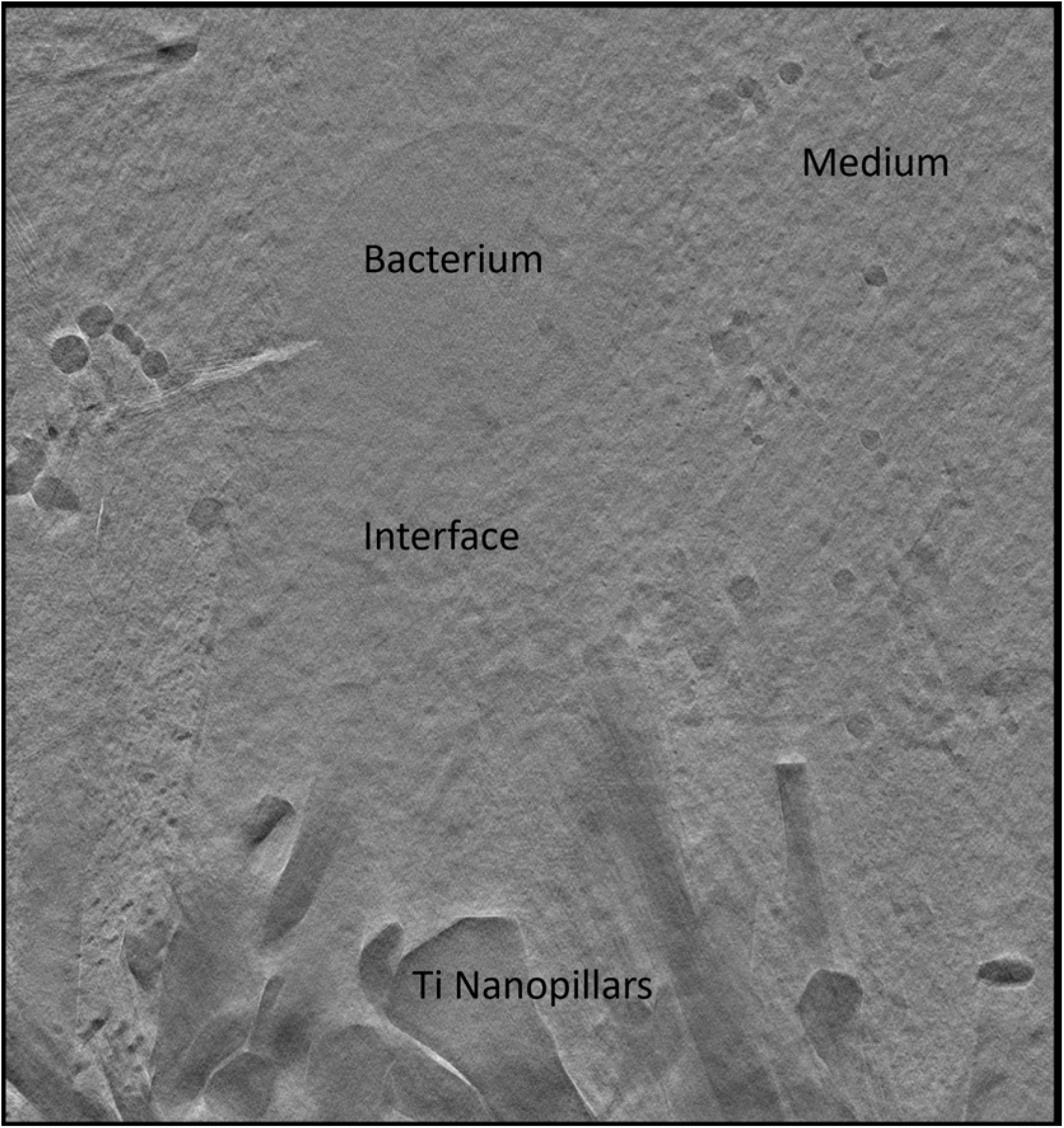
Cryo TEM tomogram showing nanopillar-bacteria interface in near-native hydrated state.

## Discussion

The main aim of this study was to establish a correlative cryo-microscopy workflow (**Figure 3-8**) enabling targeted extraction and direct structural characterisation of bacteria-nanopillar interfaces under fully hydrated conditions – addressing a long-standing methodological gap in the field. The cryo-ET data presented here serve as a proof-of-principle demonstration of this capability rather than as evidence for or against specific bactericidal mechanisms.

## Workflow capabilities and future applications

This study successfully established a correlative cryo-ET workflow for targeting a specific bacterium interacting with a nanopillar surface on clinically relevant titanium substrate under near-native hydrated conditions. Compared to conventional room-temperature preparation^[1]^ or cryo-ET on grid-adhered cells, this substrate-based *in situ* workflow avoids the need for further manipulation of cells (e.g. removal of cells or nanopillars from the original contact point) or use of chemical fixation that alters the bacterial cell wall. This approach can therefore faithfully preserve the bacteria-material interface and enable its analysis at nanoscale resolution. Despite the challenges of working with non-standard cryo-EM substrates and across geographically separated cryo-microscopy facilities, we demonstrate that high-resolution structural analysis of the bacteria-material interface is achievable with appropriate methodological adaptations. Key achievements include substrate optimisation, fiducial-based correlation, hybrid FIB strategy and proof-of-concept tomograms, validating the complete workflow from vitrification through to tomographic reconstruction.

The described workflow can be fully replicated using commercially available cryo-infrastructure and can be adapted to a range of bacterial strains and surface topographies. While HPF is preferred for optimal sample thickness and hydration, the workflow is also compatible with plunge freezing for thinner specimens. However, this depends on the thermal properties of the substrate used. The modularity of the workflow, including cryo-compatible shuttles, transfer stations and correlation strategies, allows for integration into existing cryo-EM pipelines.

The dataset presented is from sample going through the complete workflow, hence is intended as a methodological proof-of-concept and captures a vitrified snapshot of the interface at the moment of freezing. It is not intended, therefore, to fully resolve the temporal dynamics of bacterial adhesion or membrane deformation. Similarly, phenomena that depend on the presence of air-liquid interfaces, such as hypotheses involving trapped air pockets between nanostructures and bacterial surfaces, are unlikely to be preserved during HPF freezing and therefore fall outside the scope of this present approach. Nevertheless, establishing the ability to directly visualise bacteria–nanostructure interfaces under vitrified conditions opens new opportunities for investigating long-standing questions regarding the bactericidal action of bioinspired nanopatterned surfaces. Future studies building upon this workflow could systematically examine how bacterial cells interact with nanostructures across different stages of adhesion and membrane deformation, potentially enabling correlation of structural states with previously reported observations from techniques such as fluorescence microscopy, atomic force microscopy or computational simulations. In particular, combining targeted cryo-ET with larger datasets or time-resolved experimental strategies may allow investigation of processes such as initial cell wall contact, membrane deformation, and possible puncturing events at nanostructured interfaces. Furthermore, integration with complementary techniques such as fast live-cell fluorescence or ToF-SIMS imaging may help correlate kinetic observations with dynamic cellular responses. Establishing the present workflow therefore represents an important methodological step forward towards resolving the mechanistic basis of bactericidal nanotopographies and can serve as a platform for future investigations of bacterial interactions with other complex biomaterial interfaces.

## Workflow success metrics and limitations

While the workflow proved technically feasible, reproducibility remained sample- and operator-dependent, with notable challenges in milling efficiency, lift-out reliability and precision of correlation. Ice contamination during multi-facility transport and electrostatic issues during lamella transfer represented significant bottlenecks affecting overall yield. Every remount and transfer step risked the introduction of ice contamination or total loss of the sample due to manual handling. Thus, while the approach is scientifically valid, these limitations underscore the need for further instrumental and procedural refinements to achieve routine, high-throughput applications. The following sections discuss the strategic decisions, technical considerations, and future directions that have emerged from this work, paving the way for exciting new developments in the field.

### Substrate design and preparation strategy for correlative cryo-imaging

The choice of substrate material for developing this workflow involves a critical trade-off between cryo-vitrification efficiency and clinical translatability. In the cryo-EM field, sapphire disks have emerged as the gold standard for HPF due to their exceptional thermal conductivity (23-40 W/m.K), enabling rapid heat extraction and consistent vitrification of biological samples.^25–27^ In addition, they are optically transparent, allowing for easy screening in transmitted light microscopy. However, sapphire is not a material used in load-bearing medical implants due to its limited fracture toughness. Moreover, creating biomimetic nanopillar arrays on sapphire requires specialised techniques and is less established. The other potential substrate is silicon, which is widely used to study bactericidal mechanisms of nanotopographies. It can provide a flat surface enabling precise topographical control, but has fundamental limitations for medical device applications due to its low strength and fracture toughness. In our preliminary studies, after vitrification, Si substrates showed bubbling on the surface, making them unsuitable for further studies. On the other hand, titanium and its alloys (Ti-6Al-4V) represent the gold standard for orthopaedic and dental implants due to their superior biocompatibility, corrosion resistance and favourable mechanical properties, and Ti/Ti-6Al-4V nanopillar substrates are well characterised for their bactericidal action^[2]^. However, the primary disadvantage of titanium for cryo-EM workflows is its relatively lower thermal conductivity (21.9 W/m.K) compared to sapphire. Our substrate selection strategy prioritised clinical relevance over established cryo-optimised substrates, representing a broader trend in cryo-microscopy as the field expands beyond traditional cell biology applications into materials science and bioengineering. While sapphire remains optimal for fundamental cell biology studies where substrate material is irrelevant, emerging applications in biomaterials-cell characterisation increasingly requires the study of cells on clinically relevant substrates, despite their sub-optimal cryo properties.

While our previous nanopillar fabrication work utilised bulk titanium substrates of 1 mm thickness and 1 cm in diameter, the current correlative cryo-microscopy workflow required adaptation to thin substrates for two critical reasons: 1) Thermal considerations – The high thermal mass and heat capacity of bulk titanium impede the rapid cooling rates required (>10^4^ K/s) for successful vitrification. Thick metal substrates can act as heat sinks, promoting cubic ice crystal formation rather than vitreous ice, which introduces artefacts during vitrification; 2) Compatibility with cryo storage and transfers - Cryo-workflows require standardised sample dimensions to accommodate specialised transfer systems, storage containers, and sample holders across multiple instruments. Most commercial cryo-handling systems, including the Leica EM VCM, Leica EM VCT500 and associated stage holders, are optimized for 3 mm diameter planchets, matching the standard TEM grid format (3.05 mm diameter copper grids). This standardisation enabled use of commercially available cryo-storage boxes and handling tools and easy transfer between modalities without need of custom adaptors.

Our workflow demonstrates that with appropriate sample geometry modifications and advanced vitrification methods, it is possible to achieve adequate ice quality on materials like titanium.

### Vitrification method comparison

Rapid freezing of biological specimens can transform water into vitreous (amorphous) ice, effectively halting molecular motion and preserving cellular and macromolecular structures in a near-native state. This technique enables structural conservation of samples such as protein solutions, viruses, bacteria, and eukaryotic cells for EM, enabling structural analysis at resolutions approaching the atomic scale.^28^ To prevent the formation of crystalline ice, cryo-fixation must achieve extremely rapid cooling rates, typically exceeding 100,000 °C per second. Importantly, vitrified samples are stable under the high-vacuum conditions used in cryo-EM and are also compatible with cryogenic fluorescence light microscopy (cryo-fLM).^2^ While vitrification is well established for suspensions or adherent cells on EM grids, our study introduced a challenge to directly vitrify bacterial cells interacting with a titanium substrate. Titanium poses unique challenges due to its lower thermal conductivity relative to Cu and sapphire disks, size, surface topography, and optical reflectivity, which complicate both freezing efficiency and correlative imaging.

We tested both plunge freezing and HPF in this workflow. Some of the plunge frozen samples showed ice crystal formation due to poor blotting, which were discarded. Unlike perforated TEM grids used in plunge freezing, our Ti substrate was solid without grid openings, which made back side blotting inefficient. All plunge frozen samples had a crack in the ice where they were held during the plunge freezing. All samples also required clipping to an autogrid specimen cartridge to prevent damage and flake-off during the multiple handling steps required in the workflow, so that the sample could be held by the cartridge instead of the sample itself. It is important to note that some clipped samples popped out while handling in liquid nitrogen, as the c-clip could not hold the titanium substrate well enough in the cartridges. Once this occurred, they were hard to remount. These c-clips are designed to carry much thinner Cu TEM grids, which are 15-25 µm in thickness; hence, this is not a very favourable setup. Without cartridges, the plunge frozen samples were difficult to manipulate in the workflow. If thinner titanium foils were used, or much stronger c-clips are available, this popping out could have been avoided. However, we managed to locate bacteria and extract the lamella from a plunge frozen sample but at the final stage, during TEM tomography acquisition, the lamella was lost during transfer to the TEM, hence we could not acquire the cryo-tomogram from a plunge frozen sample.

With the HPF approach, a vitreous, reasonably flat substrate was achieved. The vitreous ice layer was thicker than for the plunge frozen sample, so more material had to be removed, and time taken to reach the titanium substrate during the FIB milling step. Given the large interaction volume and lack of blotting involved, vitrification was reproducible. Although we preferred HPF for our application, either of the techniques would work for sample vitrification in this workflow.

### Hardware considerations in cryo-correlative tomography workflow

The successful implementation of our cryo-tomography workflow required careful consideration of hardware capabilities to ensure sample integrity and compatibility across modalities from different manufacturers at geographically separated facilities (IMG and TESCAN). Maintaining the vitreous state throughout all steps — vitrification, fluorescence imaging, FIB milling, cryo-TEM imaging — demands a dedicated infrastructure capable of sustaining cryogenic temperatures and minimising exposure to humidity. Operating across equipment from multiple manufacturers (Leica transfer systems, TESCAN plasma FIB, JEOL TEM) introduced coordination challenges, making the standardised 3 mm sample format essential for cross-system compatibility. These hardware configurations were not incidental but foundational to the workflow, enabling high-resolution structural preservation and correlative targeting across imaging modalities at two geographically separated facilities. Our workflow demonstrates that advanced correlative cryo-microscopy is achievable without requiring all instrumentation to be co-located or from a single vendor.

Temperature control is critical throughout sample manipulation, otherwise the sample will go through thermal cycles where vitreous ice will crystallise. Therefore, the sample must be maintained below -135 °C (where initial recrystallisation starts to occur) at all times. The nanomanipulator must remain below −145 °C to preserve the vitreous state during lamella handling, while the sample stage must maintain <-145°C throughout FIB-SEM procedures. Our workflow utilised Leica VCM and VCT500 transfer systems, maintaining samples at ≤-180°C during exchange of samples between the instruments. Additionally, a humidity below 50% is essential to prevent ice contamination, which we were unable to achieve.

Cryo-compatible coating capabilities were essential for depositing conductive and protective layers without exposing samples to temperature rises or contamination. The system (Leica cryo-coater integrated with VCT500) enabled platinum or gold-palladium coating to minimize charging artefacts during imaging. Prior to FIB trench milling, an organometallic platinum layer was deposited onto the cryo-sample surface using the Gas Injection System (GIS) to provide a protective cap against ion-beam damage and curtaining artefacts. During the cryo-lift-out, however, no GIS deposition was applied for lamella-to-grid attachment, as in contrast to room-temperature workflows, the lamella welding in cryogenic conditions was achieved through redeposition from the sample material itself.

### Fiducial-based fluorescence correlation for multi-facility fluorescence guided targeting of cryo-lift-out workflows

#### The challenge: Localising Bacteria on Vitrified Titanium substrates

One aim of this workflow was to assess the capacity of targeting a specific bacterium of interest to be studied, rather than picking a random bacterium in the presence of densely packed bacterial cells. However, localising and extracting a specific bacterium attached to a titanium nanopillar surface posed significant challenges during lamella preparation for cryo-ET. Bacterial cells attached to titanium nanopillars were buried 5-25 µm beneath the vitrified ice surface, making them invisible in cryo-SEM imaging. Hence, not just X-Y coordinates but also the Z depth was important. This is where correlative cryo-fluorescence microscopy was adapted as a complementary modality to identify buried bacteria. Once bound to SYTO9, bacterial DNA fluoresced green, indicating the location of the bacteria that otherwise could not be determined by cryo-SEM alone. Here we have used a single stained bacterium as a proof-of-concept as to how cryo-fluorescence can guide subsequent electron imaging to improve the efficiency and precision of downstream cryo-EM or cryo-ET workflows. This can be expanded with other staining methods to identify specific bacteria of relevance to a study. However, while X-Y coordination was relatively successful, picking Z-direction was not always successful as we could not see the bacterial cell envelope directly interacting with the nanopillars. One reason could be that with fluorescence widefield imaging we picked a bacterium very close to the surface, yet 100-200 nm could not be resolved by widefield microscopy due to the surface topography, as seen in Figure 9. Another reason could be that the tomography analysis was confined to a small region of the entire bacterium (200 nm) and another section of the bacterium could have been attached to the surface. However, gaps between nanopillars and the bacterial cell wall at the interface have already been reported in the literature and can only be studied with a volume EM approach^29^.

To overcome limitations related to finding a specific bacterium on the samples, we 1) adapted a surface feature-based correlation strategy, and 2) developed a fiducial marker-based-correlation strategy that enabled reliable localisation of bacterial targets with high spatial precision. While both approaches were similar in concept, where a surface feature is correlated to the brightfield and SEM data, the surface feature-based strategy could be used for immediate lamella preparation using Ga-FIB, whereas the fiducial-marker strategy was needed for the combined FIB strategy where no fluorescence facility was available. For the latter, bacterial cells were preselected, positions were marked on the surface, and no image registration was required at the second facility. The key aspect was to create permanent FIB-milled fiducial patterns that were 1) cross-modality visible i.e. detectable by light microscope and electron microscopy; 2) permanent and ice contamination resistant i.e. milled features survived cryogenic storage, transport, and moderate ice contamination; 3) geometry defined - square pattern geometry enabled accurate coordinate transformation and easy image registration; and 4) reproducible i.e. the standardised pattern could be replicated across samples and laboratories easily.

These fiducials, milled into the sample surface prior to fluorescence imaging (Fig. 3), enabled reliable correlation between fluorescence data and FIB-SEM images across facilities, helping to preserve spatial accuracy throughout the workflow, despite the lack of integrated instrumentation. However, it is important to note that fiducials are visible in reflectance images and in SEM, but not in the fluorescence channel. As data is collected via two different channels of the light microscope, a slight misalignment of fluorescence and brightfield channels can lead to loss of the precise location and thus no tomogram at the end of the thinning and polishing, which is a major drawback.

Despite the isolation of a single bacterium within a 10 µm slab, a critical limitation arises after the initial 10 µm-thick slab is thinned for cryotomography due to over-milling. Without integrated fluorescence guidance, the location of the bacterium cannot be re-verified during thinning of the lamella. An additional step to check fluorescence would be helpful before thinning has started. During the thinning process, re-checking the sample in a cryo-fluorescence microscope would certainly help, but additional unloading and loading would increase the overall workflow time and the risk of ice contamination, potentially leading to total sample loss during manual handling. Consequently, we relied on getting the initial targeting right by keeping the bacterium at the centre while marking our fiducials when correlating fluorescence and SEM data sets. Even with careful alignment, there was still 500-1000 nm of uncertainty from the offset between reflected light and fluorescence channels. Despite this, we managed to capture surface-associated bacteria in 2 out of 4 attempts, which we consider reasonable given all the complications and limited availability of resources.

### Current limitations and opportunities of recent instrument developments for workflow optimisation

While the current workflow demonstrates that reliable sample preservation and spatial targeting are feasible for characterising bacteria–material interactions in their native state, several instrument enhancements could significantly improve efficiency, throughput, and lamella yield. For example, sample preparation could be improved by adding cryo-protectant during the vitrification step. It was not used here, as we wanted to characterise the interface as closely as possible to native state, but adopting cryo protection would certainly improve vitrification.

A key limitation remains the time-intensive nature of gallium-based FIB milling, which constrains the number of lamellae that can be prepared in a single session. Integration of an automated cryo-stage with active cooling would enable continuous 24-hour operation of Ga-FIB-SEM without requiring daily sample unloading, thus reducing the risk of ice contamination during loading and unloading steps. The plasma FIB-SEM system used, offer substantially higher milling rates, enabling faster trenching and bulk material removal. This is especially advantageous when processing thick vitrified samples on metallic substrates like titanium, where milling depth and area are considerable. Xenon ions were particularly effective during the initial bulk-milling stage, where large volumes of hard material had to be removed under cryogenic conditions. Compared to gallium, Xe ions enabled milling at higher currents with improved beam profiles and reduced material redeposition, producing cleaner trenches and smoother cross-sections. As a result, the overall process yielded substantially faster and more reliable lamella preparation for subsequent cryo-lift-out. It is important to note that, at room temperature, milling of titanium with a Ga beam at 30 kV is more efficient compared to plasma FIB.^30, 31^ However, the redeposition we noticed made the overall workflow less efficient in Ga-FIB at cryo temperatures. However, care must be taken to minimize potential artefacts associated with plasma beams, especially during final lamella polishing, as increased surface roughness or curtaining may occur at low polishing currents, which could compromise downstream cryo-TEM imaging. In our workflow, this was mitigated by performing the final polishing step on a separate Ga-based FIB system, which — owing to its narrower beam profile at low currents — enabled gentler and more controlled fine polishing with minimal curtaining. Although a hybrid approach combining Xe and Ga FIB systems was employed in this study, future applications of the methodology should not be limited by this setup, as the entire workflow can be implemented on a single instrument, provided the operator is aware of the respective strengths and limitations of each ion source.

Recent studies have demonstrated the cryo-FM-FIB-SEM systems for targeting fluorescently labelled organelles.^32^ This would eliminate intermediate sample transfers between imaging modalities during the workflow and reduce the risk of ice contamination and improve registration accuracy between fluorescence and electron imaging, which is critical for targeting 200 nm-thick lamellae for cryo-ET. However, these systems are not widely available and were not accessible in our workflow. Instead, we employed stand-alone cryogenic modalities across multiple facilities, which introduced additional challenges compared to fully integrated platforms. These included: 1) the need for cryogenic sample transport between facilities; 2) lack of real-time fluorescence feedback, as fluorescence correlation had to be established before plasma FIB milling without the possibility of in situ verification; 3) increased risk of ice contamination due to multiple transfers; and 4) the need for reliable coordinate preservation to relocate fluorescence-identified bacteria after transport and instrument changes. While integrated systems can mitigate many of these challenges, the present workflow demonstrates that robust correlative targeting remains feasible using distributed infrastructure.

Preparing electron-transparent cross-sections of bacteria attached to titanium nanopillar substrates presented unique challenges due to the disparate physical properties of biological and metallic materials. Differences in mechanical hardness during milling and thermal behaviour during vitrification complicate sample preparation and must be carefully considered when extending this workflow to other material systems.

A further limitation arises from the reliability of cryo-lift-out and transfer steps. Lamella loss can occur due to poor attachment to the cryo-nanomanipulator or insufficient adhesion to the TEM grid, and recovery is rarely successful and a tedious task. Electrostatic build up during lift-out, particularly in poorly conductive vitrified samples or insulating substrates, can further destabilise lamellae during transfer. The use of a copper block adaptor on the nanomanipulator needle improves redeposition efficiency and stabilises lamella attachment during welding, thereby reducing detachment risk and to enhance the reproducibility in the future.^33^

A critical bottleneck in non-integrated workflows occurs after slab extraction and placement onto a TEM grid for thinning. Without integrated fluorescence imaging on the FIB systems, the precise location of the target bacterium can no longer be tracked, and thinning often results in over-milling and loss of the target. While re-checking the grid in a cryo-fluorescence microscope could potentially help, inter-system transfers increase the risk of ice contamination, require excessive time for transfer and can lead to further sample degradation through thermal cycling. These constraints highlight the importance of fiducial-based pre-marking strategies to ensure accurate targeting throughout the workflow.

Complementary approaches may further improve targeting precision in nanomater scale. For example, the incorporation of fluorescent fiducials (e.g., beads or quantum dots) directly into the sample prior to vitrification could provide optical landmarks visible across imaging modalities. However, such fiducials require careful optimisation: they must be spectrally distinct from bacterial labels, morphologically distinguishable from bacteria, and sufficiently bright without obscuring nearby biological signals. In practice, fiducial placement, intensity, and spectral properties must be balanced to enhance correlation accuracy without compromising target visibility.

The fiducial-based surface marking strategy demonstrated here (Figure 3&4) provides a robust and accessible alternative, reducing localisation uncertainty and improving the success rate of correlative cryo-lift-out workflows for bacteria. This approach is broadly applicable beyond bacteria–nanopillar systems, including: (i) non-transparent or optically challenging substrates; (ii) multi-facility cryogenic workflows requiring sample transfer; (iii) targets embedded deep within vitrified ice; (iv) systems lacking conventional EM grid landmarks; and (v) serial lift-out workflows requiring multiple lamellae from defined regions. For laboratories without access to integrated cryo-correlative platforms, this strategy offers a practical and cost-effective solution for fluorescence-guided targeting.

Future developments, including integrated fluorescence imaging within FIB-SEM systems, cryo-compatible super-resolution microscopy, and automated image correlation tools, are expected to further enhance targeting precision and workflow efficiency. Until such systems become widely available, physical fiducial markers remain an accessible and effective solution to bridge the gap between cryo-fluorescence imaging and high-resolution electron tomography.

Together, these technological improvements would streamline the correlative cryo-tomography workflow, enhancing reproducibility and enabling higher-throughput structural studies of microbial interactions at complex material interfaces.

### Conclusions

In conclusion, this study establishes a robust cryogenic workflow for investigating bacteria-nanopillar interfaces directly on metallic substrates under hydrated conditions. By combining correlative cryo-fluorescence microscopy with targeted cryo-lift-out, we enable the site-specific preparation of electron-transparent lamellae from vitrified bacteria interacting with titanium nanopillar surfaces. The workflow provides new opportunities for *in situ*, high resolution analysis of bacteria-material interactions and offers a broadly applicable platform for studying the mechanistic basis of bactericidal nanotopographies and other biointerfaces in their native hydrated state.

## Acknowledgements

This work was supported by the Engineering and Physical Sciences Research Council under UKRI Postdoc Guarantee (grant number EP/X022609/1). Access to cryo FIB-SEM and cryo TEM was supported by Czech-BioImaging project (Ministry of Education, project number LM2023050 Czech-BioImaging) at Electron Microscopy Core Facility, IMG ASCR, Prague, Czech Republic, which is supported by MEYS CR (LM2023050 Czech-BioImaging), OP RDE (CZ.02.1.01/0.0/0.0/18_046/0016045, CZ.02.1.01/0.0/0.0/16_013/0001775) and IMG grant (RVO: 68378050). The authors gratefully acknowledge the Wolfson Bioimaging Facility and the University of Bristol Workshop for their support and assistance in this work. Authors also acknowledge the Diamond Light Source (UK) Ltd for access to the Helios Hydra via experiment number BI35964, and support by Dr. David Farmer, Dr. Michael Grange and Dr. Maud Dumoux. C.D.B is currently funded by a Helmholtz recruiting initiative grant (I-044-16-01) awarded to Liane G. Benning from the GFZ. The authors thank Ufuk Borucu at University of Bristol, Dr. Benji Bateman of Central Laser Facility of UKRI Science and Technology Facilities Council, Peter Sykes of SciTech Precision Ltd, and Samuel Zachej of Tescan Group a.s for their support. The authors thank group members of Oral Microbiology and bioMEG for their support and discussions.

## Data Availability Statement

Data will be available upon request.

## Author Information and Contributions

*Chaturanga D. Bandara; Conceptualisation, acquisition of funding, fabrication of titanium substrates, and characterisation, cryo sample preparation, data analysis, wrote the first draft*.

*Dominik Pinkas Cryo sample preparation, data acquisition with cryo Ga-FIB and cryo-TEM, edited the manuscript*.

*Martina Zanova; Data acquisition with cryo plasma-FIB-SEM, edited the manuscript*.

*Martin Uher; Data acquisition with cryo plasma-FIB-SEM, edited the manuscript*.

*Judith Mantel; Cryo sample preparation, and reviewed the manuscript*.

*Bo Su; Project supervision, administration of research funding, and reviewed the manuscript*.

*Angela H. Nobbs; Project supervision and critically revised the manuscript*.

*Paul Verkade; Project supervision, cryo sample preparation and reviewed the manuscript. All authors read and agreed to the current version of the manuscript*.

## References

(1) Baulin, V. A.; Linklater, D. P.; Juodkazis, S.; Ivanova, E. P. Exploring Broad-Spectrum Antimicrobial Nanotopographies: Implications for Bactericidal, Antifungal, and Virucidal Surface Design. ACS Nano 2025, 19 (13), 12606–12625. DOI: 10.1021/acsnano.4c15671.

(2) Bhadra, C. M.; Khanh Truong, V.; Pham, V. T. H.; Al Kobaisi, M.; Seniutinas, G.; Wang, J. Y.; Juodkazis, S.; Crawford, R. J.; Ivanova, E. P. Antibacterial titanium nano-patterned arrays inspired by dragonfly wings. Scientific Reports 2015, 5, 16817, Article. DOI: 10.1038/srep16817.

(3) Elbourne, A.; Crawford, R. J.; Ivanova, E. P. Nano-structured antimicrobial surfaces: From nature to synthetic analogues. J Colloid Interface Sci 2017, 508, 603–616. DOI: 10.1016/j.jcis.2017.07.021.

(4) Hasan, J.; Crawford, R. J.; Ivanova, E. P. Antibacterial surfaces: the quest for a new generation of biomaterials. Trends Biotechnol 2013, 31 (5), 295–304. DOI: 10.1016/j.tibtech.2013.01.017.

(5) Bandara, C. D.; Singh, S.; Afara, I. O.; Tesfamichael, T.; Wolff, A.; Ostrikov, K.; Oloyede, A. Bactericidal Effects of Natural Nanotopography of Dragonfly Wing on Escherichia coli. ACS Applied Materials & Interfaces 2017, 9 (8), 6746–6760. DOI: 10.1021/acsami.6b13666.

(6) Bandara, C. D.; Ballerin, G.; Leppänen, M.; Tesfamichael, T.; Ostrikov, K.; Whitchurch, C. B. Resolving Bio-Nano Interactions of E.coli Bacteria-Dragonfly Wing Interface with Helium Ion and 3D-Structured Illumination Microscopy to Understand Bacterial Death on Nanotopography. ACS Biomaterials Science & Engineering 2020, 6 (7), 3925–3932. DOI: 10.1021/acsbiomaterials.9b01973.

(7) Ivanova, E. P.; Hasan, J.; Webb, H. K.; Truong, V. K.; Watson, G. S.; Watson, J. A.; Baulin, V. A.; Pogodin, S.; Wang, J. Y.; Tobin, M. J.;, et al. Natural bactericidal surfaces: mechanical rupture of Pseudomonas aeruginosa cells by cicada wings. Small 2012, 8 (16), 2489–2494. DOI: 10.1002/smll.201200528.

(8) Ishantha Senevirathne, S. W. M. A.; Hasan, J.; Mathew, A.; Jaggessar, A.; Yarlagadda, P. K. D. V. Trends in Bactericidal Nanostructured Surfaces: An Analytical Perspective. ACS Applied Bio Materials 2021, 4 (10), 7626–7642. DOI: 10.1021/acsabm.1c00839.

(9) Cheng, Y.; Ma, X.; Franklin, T.; Yang, R.; Moraru, C. I. Mechano-Bactericidal Surfaces: Mechanisms, Nanofabrication, and Prospects for Food Applications. Annual Review of Food Science and Technology 2023, 14 (Volume 14, 2023), 449–472. DOI: 10.1146/annurev-food-060721-022330.

(10) Catley, T. E.; Corrigan, R. M.; Parnell, A. J. Designing Effective Antimicrobial Nanostructured Surfaces: Highlighting the Lack of Consensus in the Literature. ACS Omega 2023. DOI: 10.1021/acsomega.2c08068.

(11) Ye, Z.; Chen, X.; Zhang, P.; Zhang, X.; Zhang, X.; Chen, Y.; Liu, X.; Wang, X. Bio-Inspired Micro/Nanostructured Antibacterial Surfaces: Antibacterial Mechanisms, Design Principles, and Fabrication Methods. Adv Healthc Mater 2026, 15 (11), e03902. DOI: 10.1002/adhm.202503902 From NLM Medline.

(12) Priya, S.; Malviya, R.; Srivastava, S.; Siang, T. C.; Aseeri, A. A. Bioinspired nanostructured surfaces for antimicrobial and antifouling applications. Colloid and Interface Science Communications 2026, 70, 100863. DOI: 10.1016/j.colcom.2025.100863.

(13) Modaresifar, K.; Azizian, S.; Ganjian, M.; Fratila-Apachitei, L. E.; Zadpoor, A. A. Bactericidal effects of nanopatterns: A systematic review. Acta Biomaterialia 2019, 83, 29–36. DOI: 10.1016/j.actbio.2018.09.059.

(14) Jenkins, J.; Ishak, M. I.; Eales, M.; Gholinia, A.; Kulkarni, S.; Keller, T. F.; May, P. W.; Nobbs, A. H.; Su, B. Resolving physical interactions between bacteria and nanotopographies with focused ion beam scanning electron microscopy. iScience 2021, 24 (7), 102818. DOI: 10.1016/j.isci.2021.102818.

(15) Imihami Mudiyanselage, C. C. D. B. Characterisation of the bactericidal efficacy of natural nano-topography using dragonfly wing as a model. PhD, 2017. https://eprints.qut.edu.au/106746/.

(16) Alameda, M. T.; Osorio, M. R.; Pedraz, P.; Rodríguez, I. Mechano-Dynamic Analysis of the Bactericidal Activity of Bioinspired Moth-Eye Nanopatterned Surfaces. Advanced Materials Interfaces n/a (n/a), 2200608. DOI: 10.1002/admi.202200608.

(17) Liu, X.; Ishak, M. I.; Ma, H.; Su, B.; Nobbs, A. H. Bacterial Surface Appendages Modulate the Antimicrobial Activity Induced by Nanoflake Surfaces on Titanium. Small 2024, 20 (26), e2310149. DOI: 10.1002/smll.202310149 From NLM Medline.

(18) Wang, J.; Macdonald, B.; Cho, T. H.; Repetto, T.; Sun, K.; Tuteja, A.; Dasgupta, N. P. Bioinspired Zwitterionic Nanowires with Simultaneous Biofouling Reduction and Release. Small 2024, 20 (40), e2400784. DOI: 10.1002/smll.202400784 From NLM PubMed-not-MEDLINE.

(19) Schiøtz, O. H.; Klumpe, S.; Plitzko, J. M.; Kaiser, C. J. O. Cryo-electron tomography: en route to the molecular anatomy of organisms and tissues. Biochemical Society Transactions 2024, 52 (6), 2415–2425. DOI: 10.1042/bst20240173 (accessed 10/27/2025).

(20) Rice, W. J.; Cheng, A.; Noble, A. J.; Eng, E. T.; Kim, L. Y.; Carragher, B.; Potter, C. S. Routine determination of ice thickness for cryo-EM grids. Journal of Structural Biology 2018, 204 (1), 38–44. DOI: 10.1016/j.jsb.2018.06.007.

(21) Jenkins, J.; Mantell, J.; Neal, C.; Gholinia, A.; Verkade, P.; Nobbs, A. H.; Su, B. Antibacterial effects of nanopillar surfaces are mediated by cell impedance, penetration and induction of oxidative stress. Nature Communications 2020, 11 (1), 1626. DOI: 10.1038/s41467-020-15471-x.

(22) Zachs, T.; Schertel, A.; Medeiros, J.; Weiss, G. L.; Hugener, J.; Matos, J.; Pilhofer, M. Fully automated, sequential focused ion beam milling for cryo-electron tomography. eLife 2020, 9, e52286. DOI: 10.7554/eLife.52286.

(23) Mastronarde, D. N. Automated electron microscope tomography using robust prediction of specimen movements. J Struct Biol 2005, 152 (1), 36–51. DOI: 10.1016/j.jsb.2005.07.007 From NLM Medline.

(24) Hagen, W. J. H.; Wan, W.; Briggs, J. A. G. Implementation of a cryo-electron tomography tilt-scheme optimized for high resolution subtomogram averaging. Journal of structural biology 2017, 197 (2), 191–198. DOI: 10.1016/j.jsb.2016.06.007 From NLM Medline.

(25) Reipert, S.; Fischer, I.; Wiche, G. High-pressure freezing of epithelial cells on sapphire coverslips. Journal of Microscopy 2004, 213 (1), 81–85. DOI: 10.1111/j.1365-2818.2004.01260.x.

(26) McDonald, K.; Schwarz, H.; Müller-Reichert, T.; Webb, R.; Buser, C.; Morphew, M. Chapter 28 - “Tips and Tricks” for High-Pressure Freezing of Model Systems. In Methods in Cell Biology, Müller-Reichert, T. Ed.; Vol. 96; Academic Press, 2010; pp 671–693.

(27) Studer, D.; Humbel, B. M.; Chiquet, M. Electron microscopy of high pressure frozen samples: bridging the gap between cellular ultrastructure and atomic resolution. Histochemistry and Cell Biology 2008, 130 (5), 877–889. DOI: 10.1007/s00418-008-0500-1.

(28) Matias, V. R. F.; Al-Amoudi, A.; Dubochet, J.; Beveridge, T. J. Cryo-Transmission Electron Microscopy of Frozen-Hydrated Sections of Escherichia coli and Pseudomonas aeruginosa. Journal of Bacteriology 2003, 185 (20), 6112–6118. DOI: 10.1128/jb.185.20.6112-6118.2003.

(29) Collinson, L. M.; Bosch, C.; Bullen, A.; Burden, J. J.; Carzaniga, R.; Cheng, C.; Darrow, M. C.; Fletcher, G.; Johnson, E.; Narayan, K.;, et al. Volume EM: a quiet revolution takes shape. Nature Methods 2023, 20 (6), 777–782. DOI: 10.1038/s41592-023-01861-8.

(30) Brogden, V.; Johnson, C.; Rue, C.; Graham, J.; Langworthy, K.; Golledge, S.; McMorran, B. Material Sputtering with a Multi-Ion Species Plasma Focused Ion Beam. Advances in Materials Science and Engineering 2021, 2021 (1), 8842777. DOI: 10.1155/2021/8842777.

(31) Burnett, T. L.; Kelley, R.; Winiarski, B.; Contreras, L.; Daly, M.; Gholinia, A.; Burke, M. G.; Withers, P. J. Large volume serial section tomography by Xe Plasma FIB dual beam microscopy. Ultramicroscopy 2016, 161, 119–129. DOI: 10.1016/j.ultramic.2015.11.001 From NLM.

(32) Yang, J.; Vrbovská, V.; Franke, T.; Sibert, B.; Larson, M.; Hall, A.; Rigort, A.; Mitchels, J.; Wright, E. R. Integrated Fluorescence Microscopy (iFLM) for Cryo-FIB-milling and In-situ Cryo-ET. bioRxiv 2023, 2023.2007.2011.548578. DOI: 10.1101/2023.07.11.548578.

(33) Schiøtz, H. O.; Kaiser, O. J. C.; Klumpe, S.; Morado, R. D.; Poege, M.; Schneider, J.; Beck, F.; Klebl, P. D.; Thompson, C.; Plitzko, M. J. Serial Lift-Out: sampling the molecular anatomy of whole organisms. Nature Methods 2023. DOI: 10.1038/s41592-023-02113-5.

